# Epistatic interactions can moderate the antigenic effect of substitutions in hemagglutinin of influenza H3N2 virus

**DOI:** 10.1101/506030

**Authors:** Björn F. Koel, David F. Burke, Stefan van der Vliet, Theo M. Bestebroer, Guus F. Rimmelzwaan, Albert D.M.E. Osterhaus, Derek J. Smith, Ron A.M. Fouchier

## Abstract

We previously showed that single amino acid substitutions at seven positions in hemagglutinin determined major antigenic change of influenza H3N2 virus. Here, the impact of two such substitutions was tested in eleven representative H3 hemagglutinins to investigate context-dependence effects. The antigenic effect of substitutions introduced at hemagglutinin position 145 was fully independent of the amino acid context of the representative hemagglutinins. Antigenic change caused by substitutions introduced at hemagglutinin position 155 was variable and context-dependent. Our results suggest that epistatic interactions with contextual amino acids in the hemagglutinin can moderate the magnitude of antigenic change.

---

Influenza viruses of the H3N2 subtype have been circulating in humans since 1968 and are a major cause of annual epidemics. Antibodies against the hemagglutinin (HA) surface glycoprotein can neutralize the virus and are a critical component of our immune defense against influenza viruses. However, the HA changes over time to escape from recognition by neutralizing antibodies present in the human population. The antigenic evolution of H3N2 viruses was previously mapped using hemagglutination inhibition (HI) assay data spanning a 35-year period (1). During this period, 11 genetically and antigenically distinct clusters emerged that comprise viruses of high antigenic similarity, each of which was consecutively replaced by a new cluster of antigenically distinct viruses (Fig 1A). Antigenic cluster transitions, the major antigenic changes between clusters, were subsequently shown to be predominantly caused by single amino acid substitutions on seven key positions adjacent to the HA receptor binding site (RBS) (2). Most positions were involved in cluster transitions multiple times suggesting that possibilities for antigenic change of influenza viruses are restricted (Fig. 1B and 1C).

**FIG 1:**
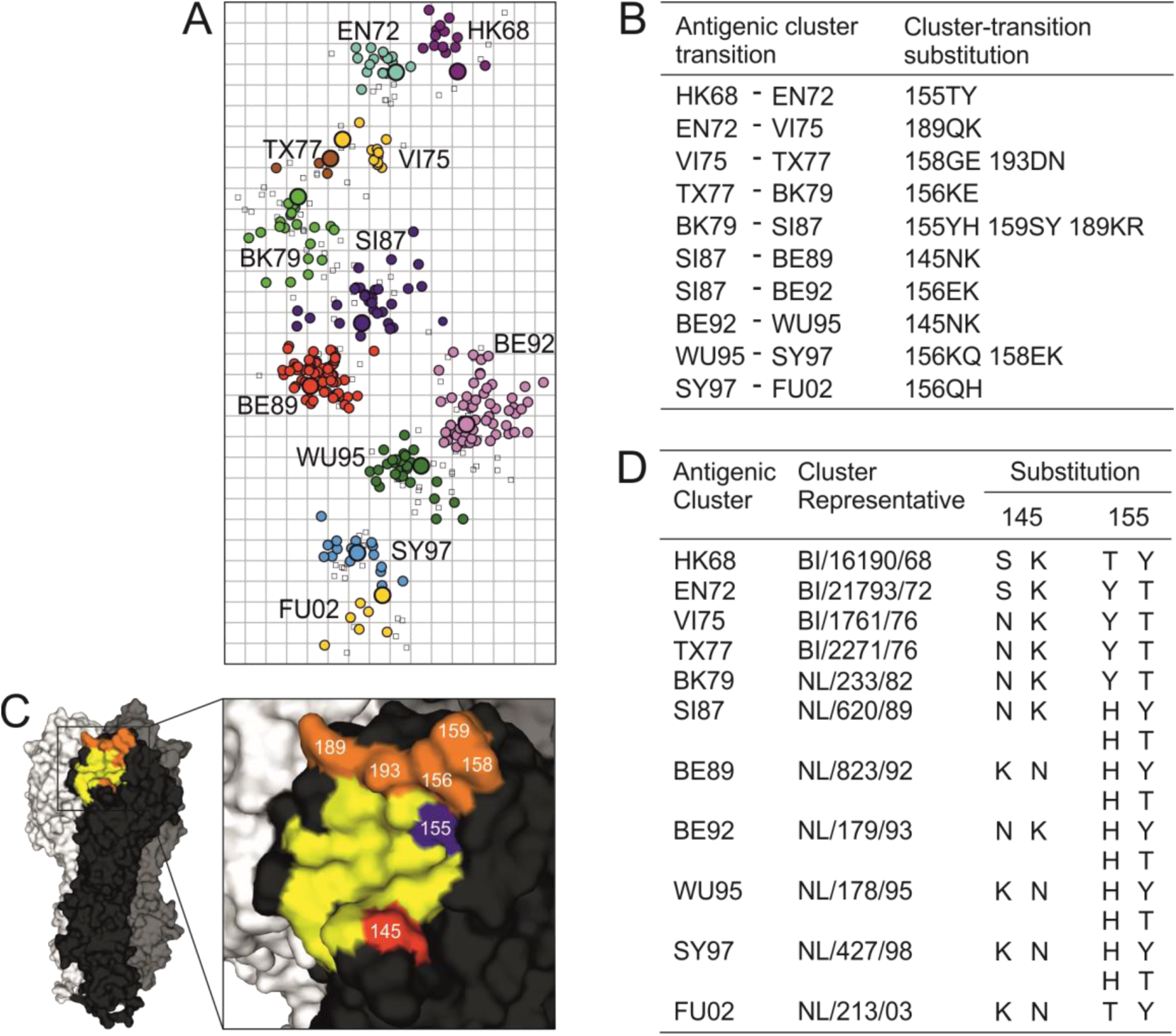
Experimental background and viruses used in this study. (A) Antigenic map of H3N2 virus antigenic evolution between 1968 and 2003. Open squares and colored circles indicate antisera and epidemic viruses, respectively. The viruses are color coded according to the antigenic cluster cluster to which the virus belongs. Both horizontal and vertical axes indicate antigenic distance, the spacing between gridlines is one antigenic unit which equals a two-fold difference in the HI assay. Letters and digits in antigenic cluster names refer to the location and year of isolation of the first vaccine strain in that cluster (HK, Hong Kong; EN, England; VI, Victoria; TX, Texas; BK, Bangkok; SI, Sichuan; BE, Beijing; WU, Wuhan; SY, Sydney; FU, Fujian). The large circles indicate the viruses used in this study to represent the antigenic cluster clusters. (B) Substitutions responsible for antigenic cluster transitions as defined in (2). (C) Amino acid positions responsible for major antigenic change during H3N2 virus antigenic evolution plotted on an A/Aichi/2/68 HA trimer (PDB accession code 5HMG). Monomers are shown in black, grey, and white, while the RBS is in yellow. Amino acid positions 145 and 155 are indicated in red and blue, respectively, while the remaining key positions are indicated in orange. (D) Mutants constructed for this study. Cluster representative viruses had the HA amino acid consensus sequence of all viruses in that cluster (described in (2)). BI, Bilthoven; NL, The Netherlands.

Epistatic interactions can shape the evolution of influenza viruses (3–6). For example, intragenic epistasis in HA has been suggested to limit the rate of antigenic evolution and to inhibit the reversal of RBS substitutions to ancestral genotypes (5–8). An important question that remains unanswered is whether the HA amino acid context in which a substitution occurs determines its ability to escape from antibody recognition.

To answer this question, we selected two substitutions that were responsible for major antigenic change during H3N2 virus antigenic evolution (Fig. 1B). Substitution 155TY was responsible for the first antigenic cluster transition of the H3N2 virus in 1972, and 155YH together with 159SY and 189KR caused an antigenic cluster transition in 1987 (2). Substitution 145NK first caused an antigenic cluster transition in 1989 after 21 years of H3N2 virus evolution in humans (1, 2). The same substitution was responsible for another cluster transition six years later. We introduced the substitutions as single mutations into the HA genes of viruses representing the 11 antigenic clusters (Fig. 1D). Depending on the amino acid at position 155 or 145, we introduced either 155Y or T, or 145K or N in the HA genes (Fig. 1D). Viruses with naturally occurring 145SN substitutions were detected from 1973 onwards (1). Between 1975 and 1989 nearly all isolated strains had 145N. However, the 145SN substitution did not contribute to major antigenic change during this period (2). When representative viruses had 145S we therefore introduced 145K, but not 145N. Substitution 155H was involved in the cluster transition that occurred in 1987 and 155H remained dominant between 1987 and 2002. For representative viruses with 155H, two modified HA genes containing either 155T or 155Y were constructed (Fig 1D). All introduced substitutions resulted in substantial changes in the biophysical properties of the amino acids. Plasmids containing wildtype or modified HA genes were used to generate recombinant viruses consisting of the (modified) HA gene and remaining genes of A/Puerto Rico/8/34 by reverse genetics (9). The presence of introduced mutations and absence of unwanted changes in HA was confirmed by Sanger sequencing. Subsequently, the antigenic properties of recombinant viruses were analyzed in HI assays using the previously defined panel of ferret antisera listed in Table S1 (2, 10). To test the antigenic difference between 155T and 155Y in representative viruses with 155H we compared the HI results of the 155T and 155Y mutants.

Mutants with substitutions at HA position 155 in HK68, EN72, VI75, TX77, SY97, and FU02 representative viruses were substantially antigenically different from their respective wildtype viruses, with up to 64-fold differences in HI titers (Fig. 2A). The 155TY amino acid difference at position 155 had a small antigenic effect in the HA context of all but one of the remaining representative viruses (SI87, BE89, BE92, WU95). Additionally, substitutions 155HT and HY had no or only modest antigenic effects in these four representative viruses— none had a more than 2-fold HI titer reduction against sera raised to viruses with homologous wildtype HAs. In the SY97 HA the 155T mutant was substantially antigenically different from the wildtype virus, but the 155Y mutant was not. Of the five representative viruses with a naturally present 155H, the 155TY amino acid difference thus had a substantial antigenic effect only in the context of a single HA. These data strongly suggest that the modest effect of the 155TY difference in multiple HAs was due to the amino acid context in which it was introduced. Thus, although the TY substitution at position 155 substantially changed the antigenic properties in more than half of the HAs tested here, the HA amino acid context in which this substitution occurs may dampen its ability to escape from antibody recognition.

**FIG 2:**
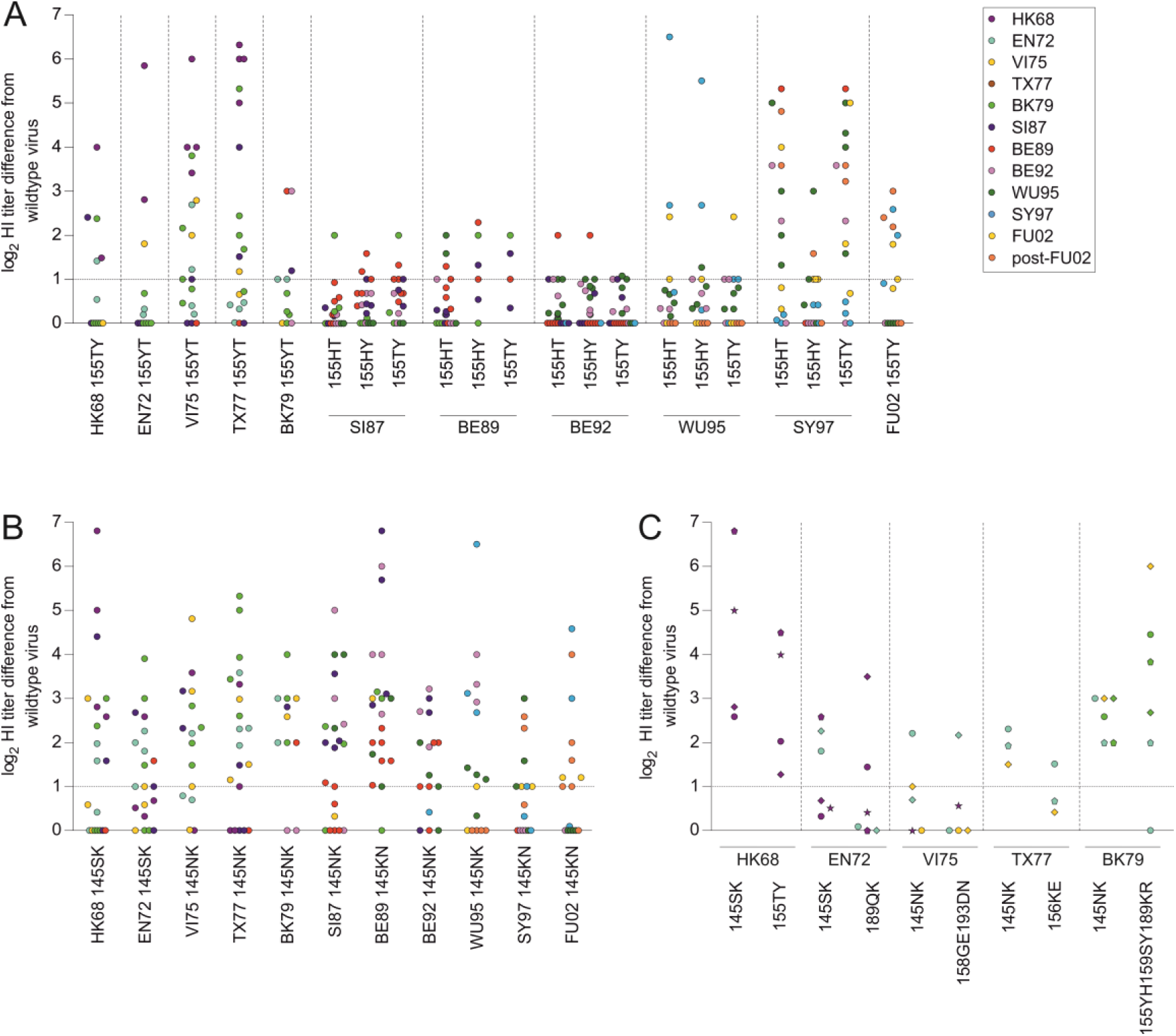
(A) HI titer differences between viruses with wild type and 155 mutant HAs. Each symbol represents the log2 HI titer difference for an individual antiserum between a representative virus and a mutant with 155TY or 155YT, or between mutants with 155HT and 155HY (indicated as 155TY for SI87, BE89, BE92, WU95, and SY97). Color coding indicates the corresponding antigenic clusters for the strains used to raise the antisera (Fig 1A). The 2-fold difference in HI titer considered to be the error of the HI assay is indicated by the dashed horizontal line. (B) HI titer differences between viruses with wildtype HA and 145K or 145N mutants. Symbols as in panel A. (C) HI titer differences between cluster representatives, 145K mutants, and cluster-transition mutants. Each symbol represents the log2 HI titer difference for an individual antiserum between viruses with wildtype and 145K mutant HA or between the wildtype and cluster-transition mutant virus. For the analysis in panel C only antisera to strains from the same or preceding antigenic clusters as the representative virus were included. Color coding as in panel A. Shapes indicate the individual antisera used for this analysis. HI data are available from Table S2.

In contrast, mutants with substitutions on position 145 of the same set of representative HAs were each antigenically distinct from their respective wildtype viruses (Fig. 2B). Thus, the magnitude of antigenic change caused by 145 NK or KN substitutions appears not to be affected by the HA amino acid context.

Substitution 145NK was first observed when it caused the antigenic cluster transition from the SI87 to the BE89 cluster (Fig. 1A and 1B). When 145K was introduced in the HA of viruses representing the antigenic clusters that circulated prior to the SI87 cluster (HK68, EN72, VI75, TX77, and BK79), it caused similar escape from inhibition by antisera to contemporary or previously circulating strains as did 145K in the SI87 representative virus (Fig. 2B). We therefore next compared the magnitude of antigenic escape by 145K to that of the cluster-transition substitutions that occurred naturally before 1989 (Fig. 1). In this analysis, only antisera to strains from the same or previous antigenic clusters as the representative virus were included, thus testing escape from antibodies induced to previously circulating strains. The magnitude of the antigenic differences caused by 145K were similar to those caused by the naturally occurring substitutions that were responsible for antigenic cluster transitions before 1989 (Fig. 2C). Thus, if viruses with 145K had appeared before the BE89 antigenic cluster they may have been sufficiently antigenically different from earlier H3N2 viruses to provide escape from population immunity.

The central question addressed in this study was if the antigenic effect of substitutions in HA is dependent on the amino acid context in which they occur. We answered this question using two substitutions known to be responsible for escape from population immunity in the past and the same analysis methods that were used to determine the contribution of these substitutions to antigenic evolution (2). The data generated for this study reflect the ability of the test viruses to escape from binding by antibodies in polyclonal ferret antisera at a fixed HA-activity. The magnitude of antigenic change caused by the introduced substitutions in the representative hemagglutinins far exceeds the antigenic change observed in studies using hemagglutinins with different binding avidities (11-13). Additionally, the large titer differences observed between sera tested to the same virus, up to 6.8 log2 HI titer differences between sera, suggest that our results are not simply a reflection of the small differences that are the resultant of variations in receptor avidity.

HA amino acid context did not affect the magnitude of antigenic change caused by substitutions introduced on position 145, nor of the majority of substitutions introduced at position 155. Thus, the ability to cause antibody escape in the HAs tested here was largely independent of the amino acid context. While these results are in agreement with the recurrent use of seven key positions for major antigenic change during influenza H3N2 virus evolution and emphasize the potential importance of these key positions for future antigenic change (2), the data also suggest that epistatic interactions govern the antigenic effect caused by the substitutions introduced HA position 155.

The differences in magnitude of the antigenic effects of 155T and 155Y substitutions versus the context-independent antigenic changes caused by the 145N and 145K substitutions in different amino acid backgrounds may be due to differences in local HA structure (Fig. 3). Position 155 is located in the depression between the 190-helix that contains conserved position 195Y and a loop that contains conserved position 153W, which are fundamental components of the RBS (14, 15). In contrast, position 145 is located on a protruding loop that may have fewer structural constraints. Therefore, the substitutions introduced at position 145 may possibly have a larger impact on the local HA structure than the substitutions introduced at position 155, resulting in the more pronounced antigenic changes in the mutants with a substitution at position 145 observed here.

**FIG 3:**
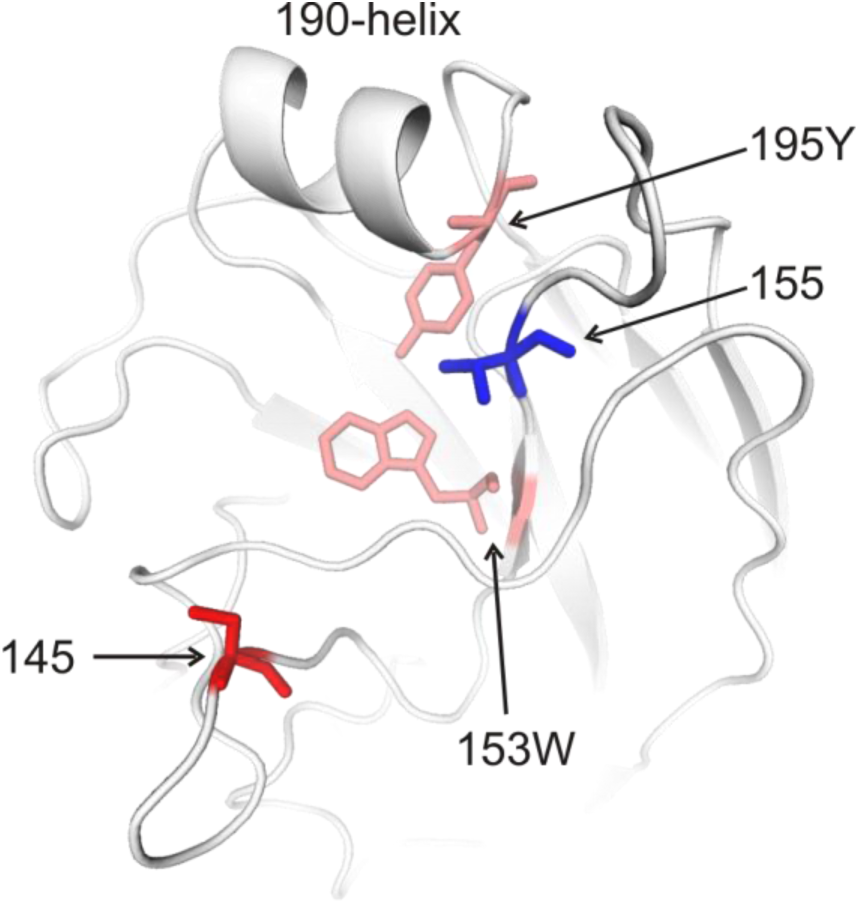
Cartoon representation of the A/Aichi/2/68 RBS area. Positions 155 and 145 are indicated in blue and red, respectively. Positions 195Y and 153W, which are conserved among influenza A virus subtypes (14, 15), are indicated in pink.

The ability of a new antigenic variant to escape from population immunity depends on the distance between the antigenic variant and the contemporary epidemic virus, which should be sufficiently large to escape from neutralization by antibodies to currently circulating viruses. Additionally, the direction of antigenic evolution should be away from all previously circulating viruses to maximize escape from recognition by antibodies to viruses responsible for earlier epidemics. We have here focused on testing the magnitude of antigenic change caused by substitutions in different HA amino acid contexts because limitations inherent to our experimental setup preclude meaningful analysis of directionality. Although our data indicate that the magnitude of antigenic change by 145K in HAs representative of early evolution strains equals that of the naturally selected escape mutants, we can make no claims about the directionality of antigenic change. Earlier work showed that epistatic interactions can affect the directionality of antigenic evolution because introduction of co-occurring mutations with cluster-transition substitutions changed the directionality of the mutant viruses without adding to the antigenic distance (2). Although many evolutionary variables may determine which viruses eventually cause an epidemic, viruses with naturally selected escape mutations perhaps had a more favorable direction of antigenic evolution compared to viruses with 145K in HAs prior to 1989, which could explain why 145K escape mutants did not emerge during the first decades of H3N2 virus evolution.

In summary, the requirement that substitutions occur in an HA context that is permissive for the protein changes that induce antibody escape suggests that the magnitude of antigenic change depends on epistatic interactions. Understanding the role of epistasis in antigenic evolution will help to evaluate the epidemic potential of newly emerging viruses.

## Acknowledgments

This work was supported by a ZonMW VICI grant and NIH contracts HHSN266200700010C and HHSN272201400008C, NIH Director’s Pioneer Award DP1-OD000490-01, European Union FP7 program EMPERIE (223498), European Union FP7 program ANTIGONE (278976), and program grant P0050/2008 from the Human Frontier Science Program.

The authors gratefully thank J.C. de Jong, B.M. Laksono, N.S. Lewis, and C.A. Russell for stimulating discussions and technical assistance.

