## Supplemental Table 1 for "Epistatic interactions can moderate the antigenic effect of substitutions in hemagglutinin of influenza H3N2 virus"

| Antigenic cluster | Antisera | | |
|---|---|---|---|
| HK68 | A/Hong Kong/1/68 | A/Hong Kong/1A/68 | A/Hong Kong/107/71 |
| | A/Hong Kong/107A/71 | | |
| EN72 | A/Bilthoven/21793/72 | A/Bilthoven/21793A/72 | A/Port Chalmers/1A/73 |
| | A/Port Chalmers/1B/73 | | |
| VI75 | A/Victoria/3A/75 | A/Victoria/3D/75 | |
| TX77 | - | | |
| BK79 | A/Netherlands/209/80 | A/Netherlands/233/82 | A/Netherlands/241/82 |
| | A/Philippines/2/82 | A/Stockholm/10A/85 | A/Stockholm/10B/85 |
| SI87 | A/Sichuan/2/87 | A/Shanghai/11A/87 | A/Shanghai/11B/87 |
| | A/Hong Kong/34A/90 | | |
| BE89 | A/Beijing/353C/89 | A/Victoria/2/90 | A/Victoria/2B/90 |
| | A/Lyon/1149/91 | A/Lyon/1149A/91 | A/Paris/548B/92 |
| | A/Netherlands/823B/92 | | |
| BE92 | A/Beijing/32/92 | A/Shangdong/9/93 | A/Shangdong/9B/93 |
| | A/Johannesburg/33A/94 | | |
| WU95 | A/Lyon/2279A/95 | A/Lyon/2279B/95 | A/Nanchang/933/95 |
| | A/Wuhan/359B/95 | A/Brisbane/8/96 | A/Brisbane/8B/96 |
| SY97 | A/Sydney/5A/97 | A/Netherlands/118/01 | A/Netherlands/88/03 |
| FU02 | A/Fujian/411/02 | A/Netherlands/22/03 | A/Wellington/001/04 |
| post-FU02 | A/Hiroshima/052/05 | A/Wisconsin/67/05 | A/Netherlands/42/06 |
| | A/Brisbane/010/07 | A/Perth/016/09 | |

Table S1 Ferret antisera used in this study. Ferret antisera were raised as described in Koel *et al.*, Science, 2013, 342:976–9. Antisera raised to the same strain are annotated A, B, C, etcetera in addition to the strain name. For example, antisera A/Hong Kong/1/68 and A/Hong Kong/1A/68 were raised to the same strain. The data obtained using antisera raised to the same strain were analyzed as individual antisera and were not averaged.
