## Supplemental Table 2 for "Epistatic interactions can moderate the antigenic effect of substitutions in hemagglutinin of influenza H3N2 virus"

| | HK/1/68 | HK/1A/68 | HK/107/71 | HK/107A/71 | BI/21793A/72 | BI/21793/72 | PC/1A/73 | PC/1B/73 | VI/3A/75 | VI/3D/75 | NL/209B/80 | NL/209/80 | NL/233/82 | NL/241/82 | PH/2/82 | ST/10A/85 | ST/10B/85 |
|---|---|---|---|---|---|---|---|---|---|---|---|---|---|---|---|---|---|
| HK68 | 1920 | 4480 | 560 | 2560 | 480 | 160 | 120 | 110 | 120 | 40 | <10 | * | <10 | <20 | <10 | 26 | 40 |
| HK68 145SK | 320 | 640 | <10 | 80 | 160 | 160 | 160 | 28 | 80 | <10 | <10 | * | <10 | <20 | <10 | <10 | <10 |
| HK68 155TY | * | 1600 | * | 160 | 1280 | * | * | 160 | * | 40 | <10 | * | <10 | <20 | <10 | <10 | * |
| EN72 | 100 | 320 | 30 | 100 | 2240 | 960 | 320 | 320 | 240 | 140 | 25 | * | <10 | <20 | <10 | <10 | <10 |
| EN72 145SK | 80 | 200 | <10 | 70 | 640 | 200 | 160 | 80 | 160 | 70 | 25 | * | 14 | <20 | <10 | 75 | 40 |
| EN72 155YT | * | 2240 | * | 5760 | 2560 | * | * | 400 | * | 40 | 40 | * | <10 | <20 | <10 | <10 | * |
| VI75 | 10 | <10 | <10 | 60 | 554 | 259 | 120 | 171 | 1109 | 1280 | 716 | 560 | 277 | 140 | 320 | 381 | 240 |
| VI75 145NK | * | 60 | * | 60 | 120 | 160 | * | 99 | 1120 | 640 | 80 | <40 | 70 | 50 | 45 | 75 | * |
| VI75 155YT | 160 | 80 | 320 | 640 | 640 | 40 | 280 | 226 | 160 | 320 | 160 | * | * | <20 | 160 | 277 | 140 |
| TX77 | <30 | <10 | 10 | 50 | 396 | 60 | 200 | 153 | 126 | 905 | 3168 | 1120 | 1224 | 400 | 434 | 914 | 1600 |
| TX77 145NK | <30 | <10 | <10 | <10 | 80 | <10 | 40 | 40 | 280 | 320 | 400 | * | 80 | <20 | 40 | 150 | 50 |
| TX77 155YT | 480 | 400 | 640 | 3200 | 400 | 80 | 160 | 212 | 80 | 400 | 985 | * | * | <20 | 80 | 554 | 400 |
| BK79 | * | * | * | * | 40 | * | * | 20 | * | 320 | 3840 | * | 2560 | * | 1280 | 1280 | * |
| BK79 145NK | * | * | * | * | <10 | * | * | <10 | * | 40 | 640 | * | 160 | * | 320 | 160 | * |
| BK79 155YT | * | * | * | * | 80 | * | * | 40 | * | 320 | 3200 | * | 1600 | * | 1120 | 1280 | * |
| SI87 | 80 | 160 | <10 | 25 | <10 | <10 | <10 | <10 | <10 | <10 | <40 | <40 | <10 | <20 | 24 | 458 | 1568 |
| SI87 145NK | * | * | * | * | * | * | * | * | * | * | 25 | * | * | <20 | * | 89 | 400 |
| SI87 155HT | * | * | * | * | * | * | * | * | * | * | <10 | * | * | <20 | 20 | 358 | * |
| SI87 155HY | * | * | * | * | * | * | * | * | * | * | 20 | * | * | <20 | * | 423 | 1600 |
| BE89 | * | * | * | * | * | * | * | * | * | * | <10 | * | * | <20 | <10 | <20 | 80 |
| BE89 145KN | * | * | * | * | * | * | * | * | * | * | 40 | * | * | <20 | * | 89 | 640 |
| BE89 155HT | * | * | * | * | * | * | * | * | * | * | <10 | * | * | <20 | * | <20 | 70 |
| BE89 155HY | * | * | * | * | * | * | * | * | * | * | * | * | * | * | <10 | 40 | * |
| BE92 | * | * | * | * | <10 | * | * | <10 | * | <10 | <10 | * | <20 | * | <10 | <10 | 35 |
| BE92 145NK | * | * | * | * | * | * | * | * | * | * | * | * | * | * | * | * | * |
| BE92 155HT | * | * | * | * | * | * | * | * | * | * | * | * | * | * | * | * | * |
| BE92 155HY | * | * | * | * | * | * | * | * | * | * | * | * | * | * | * | * | * |
| WU95 | * | * | * | * | * | * | * | * | * | * | * | * | * | * | * | * | * |
| WU95 145NK | * | * | * | * | * | * | * | * | * | * | * | * | * | * | * | * | * |
| WU95 155HT | * | * | * | * | * | * | * | * | * | * | * | * | * | * | * | * | * |
| WU95 155HY | * | * | * | * | * | * | * | * | * | * | * | * | * | * | * | * | * |
| SY97 | * | * | * | * | * | * | * | * | * | * | * | * | * | * | * | * | * |
| SY97 145KN | * | * | * | * | * | * | * | * | * | * | * | * | * | * | * | * | * |
| SY97 155HT | * | * | * | * | * | * | * | * | * | * | * | * | * | * | * | * | * |
| SY97 155HY | * | * | * | * | * | * | * | * | * | * | * | * | * | * | * | * | * |
| FU02 | * | * | * | * | * | * | * | * | * | * | * | * | * | * | * | * | * |
| FU02 145KN | * | * | * | * | * | * | * | * | * | * | * | * | * | * | * | * | * |
| FU02 155TY | * | * | * | * | * | * | * | * | * | * | * | * | * | * | * | * | * |

| | SH/11A/87 | SH/11B/87 | SI/2/87 | BE/353C/89 | HK/34A/90 | VI/2/90 | VI/2B/90 | LY/1149/91 | LY/1149A/91 | BE/32/92 | NL/823B/92 | PA/548B/92 | SD/9/93 | SD/9B/93 | JO/33A/94 | LY/2279A/95 |
|---|---|---|---|---|---|---|---|---|---|---|---|---|---|---|---|---|
| HK68 | <10 | 15 | 106 | * | * | * | * | <10 | * | * | * | * | * | * | * | * |
| HK68 145SK | <10 | <10 | <10 | * | * | * | * | <10 | * | * | * | * | * | * | * | * |
| HK68 155TY | <10 | * | 20 | * | * | * | * | * | * | * | * | * | * | * | * | * |
| EN72 | <10 | <10 | <10 | * | * | * | * | <10 | * | * | * | * | * | * | * | * |
| EN72 145SK | <10 | 10 | 32 | * | * | * | * | 15 | * | * | * | * | * | * | * | * |
| EN72 155YT | <10 | * | <10 | * | * | * | * | * | * | * | * | * | * | * | * | * |
| VI75 | <10 | <10 | <10 | * | * | * | * | <10 | <10 | <10 | * | * | * | <10 | * | * |
| VI75 145NK | 25 | * | 45 | * | * | * | * | * | * | * | * | * | * | * | * | * |
| VI75 155YT | <10 | <10 | <20 | * | * | * | * | <10 | * | * | * | * | * | * | * | * |
| TX77 | <10 | <10 | 14 | * | * | * | * | <10 | <10 | 20 | * | * | * | <10 | * | * |
| TX77 145NK | <10 | <10 | <10 | * | * | * | * | <10 | * | * | * | * | * | * | * | * |
| TX77 155YT | <10 | 80 | 40 | * | * | * | * | <10 | * | * | * | * | * | * | * | * |
| BK79 | * | * | 35 | * | * | * | * | * | <10 | <10 | * | * | * | <10 | * | * |
| BK79 145NK | * | * | <10 | * | * | * | * | * | 20 | <10 | * | * | * | <10 | * | * |
| BK79 155YT | * | * | 80 | * | * | * | * | * | 40 | 40 | * | * | * | <10 | * | * |
| SI87 | 988 | 589 | 887 | 240 | 320 | 560 | 1280 | 421 | 226 | 160 | 320 | 320 | 80 | 40 | <10 | <10 |
| SI87 145NK | 240 | 160 | 75 | 240 | 80 | * | 1280 | 640 | 480 | <10 | 320 | 640 | 15 | <10 | <10 | 25 |
| SI87 155HT | 1109 | * | 1131 | 160 | 320 | 640 | 1280 | 800 | 160 | 140 | 280 | 320 | * | 40 | <10 | * |
| SI87 155HY | 847 | 800 | 669 | 80 | 160 | * | 640 | 320 | 100 | 120 | 200 | 200 | 50 | 25 | <10 | <10 |
| BE89 | 55 | 65 | 12 | 80 | <10 | 800 | 640 | 784 | 640 | 20 | 640 | 1280 | <32 | <10 | <10 | <40 |
| BE89 145KN | 396 | 560 | 620 | 320 | 560 | * | 1920 | 1600 | 2560 | 320 | 3200 | 3840 | 100 | 320 | 80 | 40 |
| BE89 155HT | 63 | 80 | <20 | 100 | <10 | * | 640 | 1920 | 1120 | 40 | 960 | 2560 | <10 | <10 | <10 | <10 |
| BE89 155HY | 80 | * | 30 | * | * | 400 | * | 3840 | * | * | * | * | * | * | * | * |
| BE92 | <10 | <10 | 32 | 20 | 640 | * | 20 | <10 | <10 | 149 | <10 | <20 | 960 | 1041 | 1485 | 139 |
| BE92 145NK | <10 | * | <10 | <10 | 80 | * | <10 | * | 10 | 40 | 10 | <20 | * | 160 | 160 | * |
| BE92 155HT | <20 | <10 | 30 | 20 | 640 | * | <10 | <10 | <10 | 160 | <10 | <20 | 1920 | 1600 | 1280 | 160 |
| BE92 155HY | <10 | <10 | 20 | 20 | 640 | * | <10 | <10 | <10 | 80 | <10 | <20 | 1600 | 1280 | 1280 | 80 |
| WU95 | <10 | <10 | <10 | 40 | 160 | * | 20 | 40 | 80 | 32 | 20 | <20 | 320 | 160 | 254 | 1431 |
| WU95 145NK | * | * | * | * | * | * | * | * | * | 320 | * | * | * | 640 | 1920 | * |
| WU95 155HT | * | * | * | * | * | * | * | * | * | 40 | * | * | * | 160 | 320 | * |
| WU95 155HY | * | * | * | * | * | * | * | * | * | 20 | * | * | * | 80 | 320 | * |
| SY97 | * | * | * | * | * | * | * | * | <10 | <10 | * | * | * | <10 | <10 | 40 |
| SY97 145KN | * | * | * | * | * | * | * | * | <10 | <10 | * | * | * | <10 | <10 | <10 |
| SY97 155HT | * | * | * | * | * | * | * | * | 200 | 25 | * | * | * | 60 | <10 | 100 |
| SY97 155HY | * | * | * | * | * | * | * | * | <10 | <10 | * | * | * | <10 | <10 | <10 |
| FU02 | * | * | * | * | * | * | * | * | <10 | <10 | * | * | * | <10 | <10 | <10 |
| FU02 145KN | * | * | * | * | * | * | * | * | * | * | * | * | * | * | <10 | * |
| FU02 155TY | * | * | * | * | * | * | * | * | * | * | * | * | * | * | <10 | * |

| | LY/2279B/95 | NA/933/95 | WU/359B/95 | BR/8/96 | BR/8B/96 | SY/5A/97 | NL/118/01 | FU/411/02 | NL/22/03 | NL/88/03 | WE/001/04 | HI/052/05 | WN/67/05 | NL/42/06 | BR/010/07 | PE/016/09 |
|---|---|---|---|---|---|---|---|---|---|---|---|---|---|---|---|---|
| HK68 | * | * | * | * | * | * | * | * | * | * | * | * | * | * | * | * |
| HK68 145SK | * | * | * | * | * | * | * | * | * | * | * | * | * | * | * | * |
| HK68 155TY | * | * | * | * | * | * | * | * | * | * | * | * | * | * | * | * |
| EN72 | * | * | * | * | * | * | * | * | * | * | * | * | * | * | * | * |
| EN72 145SK | * | * | * | * | * | * | * | * | * | * | * | * | * | * | * | * |
| EN72 155YT | * | * | * | * | * | * | * | * | * | * | * | * | * | * | * | * |
| VI75 | * | * | * | * | * | * | * | * | * | * | * | * | * | * | * | * |
| VI75 145NK | * | * | * | * | * | * | * | * | * | * | * | * | * | * | * | * |
| VI75 155YT | * | * | * | * | * | * | * | * | * | * | * | * | * | * | * | * |
| TX77 | * | * | * | * | * | * | * | * | * | * | * | * | * | * | * | * |
| TX77 145NK | * | * | * | * | * | * | * | * | * | * | * | * | * | * | * | * |
| TX77 155YT | * | * | * | * | * | * | * | * | * | * | * | * | * | * | * | * |
| BK79 | * | * | * | * | * | * | * | * | * | * | * | * | * | * | * | * |
| BK79 145NK | * | * | * | * | * | * | * | * | * | * | * | * | * | * | * | * |
| BK79 155YT | * | * | * | * | * | * | * | * | * | * | * | * | * | * | * | * |
| SI87 | <10 | * | <10 | <30 | <10 | <30 | * | * | * | * | * | * | * | * | * | * |
| SI87 145NK | 80 | * | * | * | 80 | * | * | * | * | * | * | * | * | * | * | * |
| SI87 155HT | <10 | * | * | * | <10 | * | * | * | * | * | * | * | * | * | * | * |
| SI87 155HY | <10 | * | * | * | <10 | * | * | * | * | * | * | * | * | * | * | * |
| BE89 | <10 | * | 40 | <30 | 15 | <30 | * | * | * | * | * | * | * | * | * | * |
| BE89 145KN | 40 | * | * | * | 50 | * | * | * | * | * | * | * | * | * | * | * |
| BE89 155HT | <10 | * | * | * | <10 | * | * | * | * | * | * | * | * | * | * | * |
| BE89 155HY | * | * | * | * | * | * | * | * | * | * | * | * | * | * | * | * |
| BE92 | 320 | 40 | 67 | 240 | 160 | <30 | <10 | <10 | <30 | <10 | <10 | <10 | <10 | <10 | * | * |
| BE92 145NK | 640 | 40 | 160 | * | 640 | <40 | * | * | * | * | * | * | * | * | * | * |
| BE92 155HT | 240 | 20 | 57 | 280 | 320 | <30 | * | * | * | * | * | * | * | * | * | * |
| BE92 155HY | 280 | 20 | 120 | 160 | 320 | <30 | * | * | * | * | * | * | * | * | * | * |
| WU95 | 1431 | 127 | 538 | 1280 | 1542 | 130 | 453 | <10 | <30 | 32 | <10 | <10 | <10 | <10 | * | * |
| WU95 145NK | 640 | 160 | 200 | * | 640 | <30 | <10 | <20 | <30 | <10 | <10 | <10 | <10 | <10 | * | * |
| WU95 155HT | 1280 | 200 | 320 | * | 1120 | 80 | <10 | <20 | 80 | <10 | <10 | <10 | <10 | <10 | * | * |
| WU95 155HY | 800 | 160 | 400 | * | 640 | 160 | 10 | <20 | <30 | <10 | <10 | <10 | <10 | <10 | * | * |
| SY97 | * | <10 | 10 | * | 60 | 1280 | 1280 | 640 | 640 | 640 | 20 | * | <10 | 20 | * | * |
| SY97 145KN | * | <10 | <10 | * | 20 | 1280 | 1600 | 320 | 320 | 320 | 30 | * | 30 | 100 | * | * |
| SY97 155HT | * | 160 | 80 | * | 240 | 1120 | 1280 | 512 | 1120 | 672 | 320 | * | 140 | 240 | * | * |
| SY97 155HY | * | <10 | <10 | * | 80 | 960 | 1280 | 320 | 320 | 480 | <20 | * | 15 | 20 | * | * |
| FU02 | <10 | <10 | <10 | * | <10 | 640 | 599 | 1109 | 1109 | 480 | 160 | <10 | 80 | 423 | 160 | <10 |
| FU02 145KN | <10 | <10 | <10 | * | <10 | 80 | 640 | 2560 | 2560 | 20 | 320 | 80 | 160 | 1280 | 640 | <10 |
| FU02 155TY | <10 | <10 | <10 | * | <10 | 160 | 320 | 320 | 640 | 80 | 80 | <10 | 10 | 80 | 35 | <10 |
